# Oxytocin shapes spontaneous activity patterns in the developing visual cortex by activating somatostatin interneurons

**DOI:** 10.1101/866251

**Authors:** Paloma P Maldonado, Alvaro Nuno-Perez, Jan Kirchner, Elizabeth Hammock, Julijana Gjorgjieva, Christian Lohmann

## Abstract

Spontaneous network activity shapes emerging neuronal circuits during early brain development, however how neuromodulation influences this activity is not fully understood. Here, we report that the neuromodulator oxytocin powerfully shapes spontaneous activity patterns. *In vivo*, oxytocin strongly decreased the frequency and pairwise correlations of spontaneous activity events in visual cortex (V1), but not in somatosensory cortex (S1). This differential effect was a consequence of oxytocin only increasing inhibition in V1 and increasing both inhibition and excitation in S1. The increase in inhibition was mediated by the depolarization and increase in excitability of somatostatin^+^ (SST) interneurons specifically. Accordingly, silencing SST^+^ neurons pharmacogenetically fully blocked oxytocin’s effect on inhibition *in vitro* as well its effect on spontaneous activity patterns *in vivo*. Thus, oxytocin decreases the excitatory/inhibitory ratio and modulates specific features of V1 spontaneous activity patterns that are crucial for refining developing synaptic connections and sensory processing later in life.

## Introduction

In the developing brain, neuronal connections form with remarkable precision. First, axons grow to predetermined target areas guided by molecular cues. Subsequently, activity-dependent processes refine synaptic connections^1–3^: already before the senses become active, spontaneous activity drives synaptic refinement to prepare the brain for interacting with the outside world. Finally, circuits adapt to the prevalent environmental conditions through sensory experience-driven plasticity mechanisms.

Spontaneous activity is expressed in specific patterns and these patterns are crucial for the appropriate wiring of neurons. For example, in the developing retina, waves of spontaneous activity travel at specific speeds, in various directions and with different wave front shapes^4,5^. Retinal waves drive highly structured activity patterns in the central visual system including the primary visual cortex^6–8^. Perturbing these activity patterns leads to miswiring of the central visual system^9–11^.

In the adult, neuronal responses triggered by sensory stimuli are not uniform for a given stimulus, but vary depending on the internal state of an animal. Neuromodulators adapt neuronal circuits to the behavioral state of an animal to respond appropriately to stimuli when internal or environmental conditions change^12–14^. A powerful neuromodulator, oxytocin, can alter sensory perception depending on the internal state of an animal. In the mature brain, oxytocin enhances spike fidelity and reduces noise in the hippocampus^15^, increases the sensitivity of auditory cortex neurons to pup calls^16^, regulates sexual behavior in female mice^17^ and dampens fear responses by modulating neuronal activity in the amygdala^18^. Oxytocin mediates many of these effects by tuning inhibitory synaptic function or interneuron excitability.

Although, oxytocin exerts powerful behavioral control in adult animals, oxytocin levels and oxytocin receptor expression are actually higher in the developing neocortex compared to the adult neocortex. Oxytocin levels in the neocortex peak during the first postnatal week and decrease until the end of the third postnatal week when they reach adult levels^19^. Similarly, oxytocin receptor ligand binding, immunolabeling and mRNA expression in the cortex and other brain areas are maximal during the second postnatal week and decrease thereafter^19–21^. In line with its high expression levels oxytocin controls synaptic transmission in the developing cortex. For example, oxytocin triggers a temporary switch of GABA receptor action from excitatory to inhibitory to protect neurons against the risk of excitatory toxicity during birth by altering intracellular chloride concentrations^22,23^. In addition, oxytocin is required for cross-modal experience-driven synaptic plasticity during the first two weeks of life^19^. However, it has been unclear whether spontaneous activity patterns, which drive synaptic plasticity before experience-driven refinement occurs^24^, are regulated by oxytocin as well.

Here, we asked whether oxytocin signaling shapes neuronal activity patterns in the primary visual cortex (V1) before the eyes open during the second postnatal week and compared it with its role in the primary somatosensory cortex (S1) which is already used for passive sensing at this age. We found that oxytocin affected these sensory cortices differentially: in the visual cortex oxytocin modulated the frequency and correlation of spontaneous activity patterns strongly, but it did not change spontaneous activity patterns in S1. We found corresponding differences in oxytocin modulation of excitatory and inhibitory synaptic activity in V1 and S1. Finally, we provide evidence that oxytocin-mediated activation of somatostatin expressing (SST) interneurons controls specific characteristics of spontaneous activity patterns in V1 known to determine the refinement of synaptic connections in the visual cortex prior to eye opening.

## Results

### Oxytocin modulates cortical spontaneous network activity differentially across sensory areas

To study the role of the neuromodulator oxytocin in the developing cortex, we first asked whether oxytocin receptor activation modulates large-scale spontaneous activity patterns. We used *in utero* electroporation to express the calcium sensor GCaMP6s in layer 2/3 pyramidal cells across V1, higher visual areas and the barrel cortex. Then we performed wide-field *in vivo* calcium imaging in lightly anesthetized pups between postnatal days (P) 9 and 13. We observed spontaneous network activity in V1, S1 and higher visual areas (Fig. 1a, b). Frequently, network events were confined to individual sensory regions, but sometimes occurred across the entire field of view. After topical application of oxytocin (1 µM) onto the cortical surface, the occurrence of network events was strongly decreased in V1, but only modestly changed in S1 (Fig. 1a-c). In fact, the response of both areas differed significantly (V1: −43.2 ± 7.2%; S1: −7.9 ± 6.1%; p = 0.0024; n = 7, unpaired two-tailed t-test). The activity decrease in V1 was very pronounced during the first 10 minutes after oxytocin application and reduced, but still significant, thereafter (Fig. 1d, last 20 minute recording, p = 0.0156; paired two-tailed Wilcoxon test). Network event area, amplitude or duration were not affected (Fig. 1e-g).

**Figure 1.**
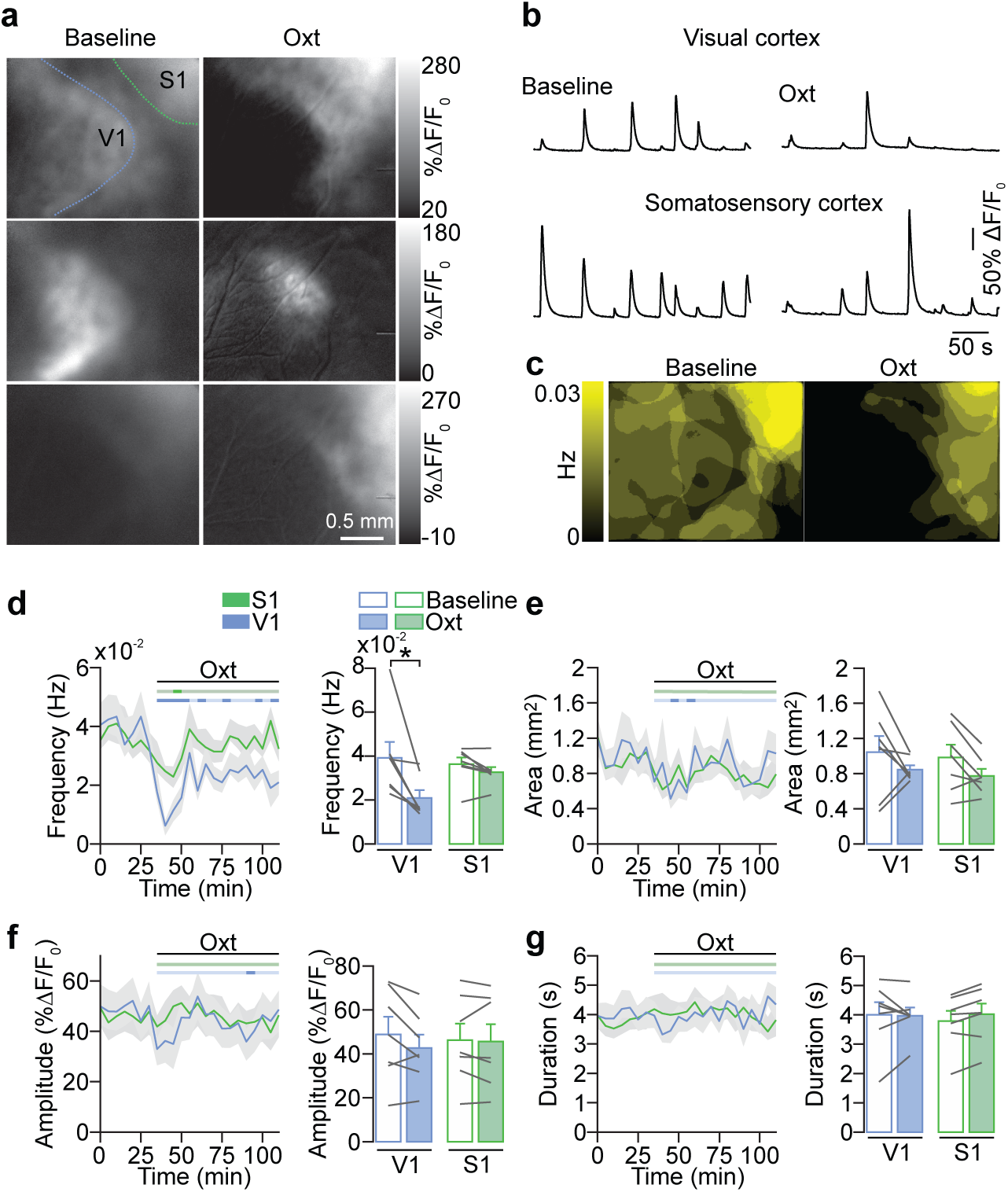
Oxytocin affects spontaneous network events differentially across sensory cortices. **a.** Wide-field calcium imaging of spontaneous activity in V1 and S1 before eye opening. Left, single frame images depicting network events activating V1 and/or S1 during baseline recordings before oxytocin application. Right, network events after oxytocin (Oxt) application. **b.** Traces show fluorescent changes in V1 and S1 before and after oxytocin application. **c.** Superimposition of all network events detected during a five-minute baseline recording (left) and after oxytocin application (right). Color code indicates the frequency of the detected events. V1 activity is strongly reduced. **d.** Network event frequency in V1 and S1 during baseline and after oxytocin application. Time courses represent five minute-averages. The horizontal bars above the line plots indicate the time points when the values for each time bin differed significantly from baseline (dark shades, paired two-tailed t-test, p < 0.05, without multi-measurement correction). *p = 0.0156 (n = 7 animals, Wilcoxon test). **e.** Network event area. **f.** Network event amplitude. **g.** Network event duration.

### Oxytocin desynchronizes spontaneous V1 network activity

To evaluate how oxytocin modulates the activity of individual neurons in the developing cortical network we performed *in vivo* two-photon calcium imaging in V1 of anesthetized neonatal mice. Layer 2/3 cells were labeled with the calcium indicator Oregon Green BAPTA-1 (OGB-1) by bolus loading^25^. Oxytocin application decreased the frequency of calcium events transiently (Fig. 2a-c) without affecting their amplitude (Fig. 2d), in line with our wide-field experiments.

**Figure 2.**
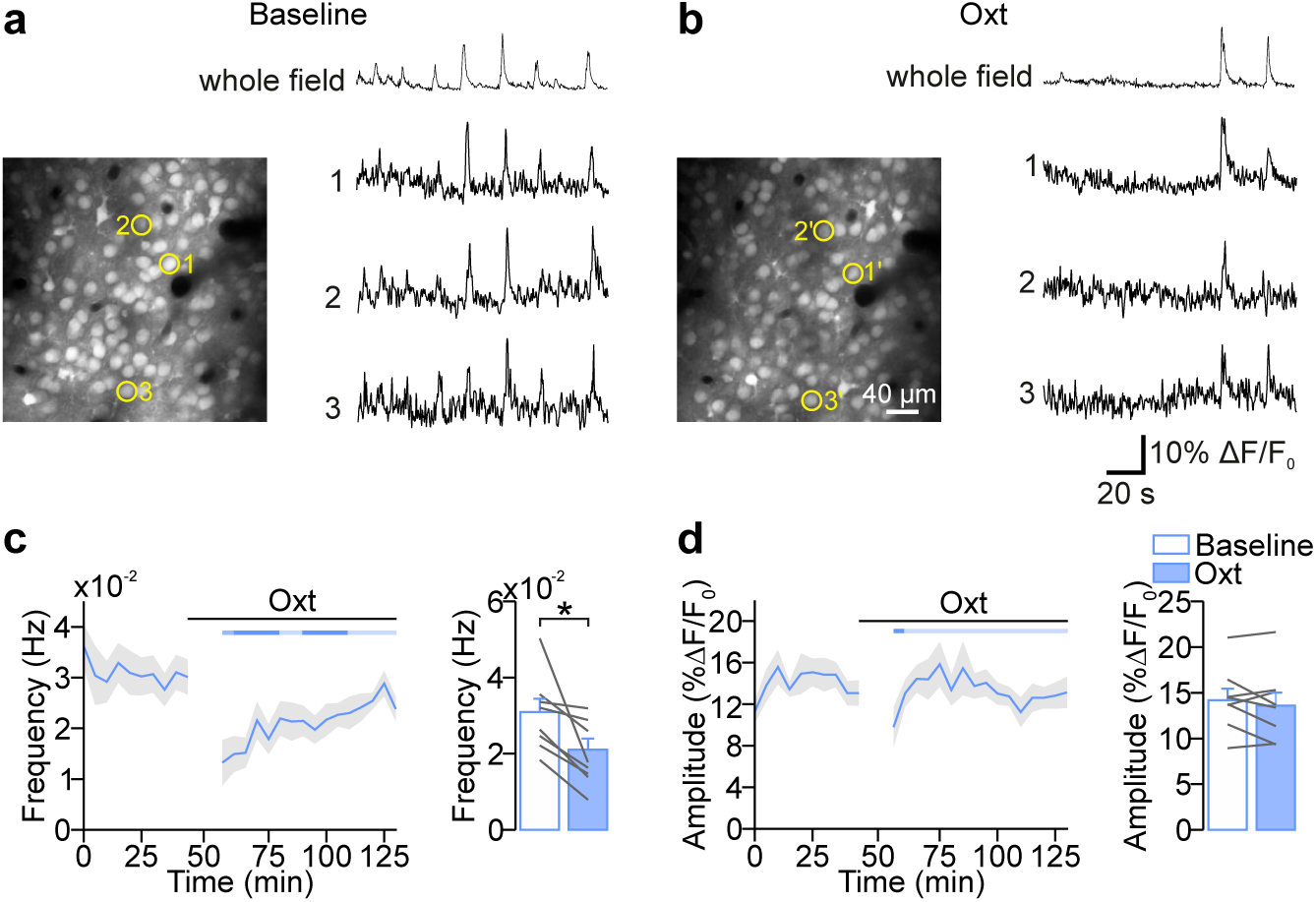
Oxytocin reduces network activity of single neurons in V1. **a, b**. V1 network activity before and after oxytocin application. Left, layer 2/3 neurons labeled with the calcium indicator Oregon Green-BAPTA 1. Traces show fluorescent changes of three example neurons and the average activity across all cells. **b.** Network event frequency during baseline and after oxytocin application. Imaging resumed approximately 10 minutes after oxytocin application. The frequency of network events was reduced after oxytocin application. The horizontal bar indicates significant deviations from baseline as in Figure 1 (dark shades, paired two-tailed t-test, p < 0.05, without multi-measurement correction). *p = 0.0154 (n = 8, paired two-tailed t-test). **d.** Network event amplitude during baseline and after oxytocin application.

Since the correlational structure of spontaneous network activity determines its role in network refinement^26^, we also investigated whether oxytocin affected the pairwise correlations between spontaneously active neurons. We observed that oxytocin application decreased the mean Pearson correlation coefficients across pairs of neurons (Fig. 3a, b). In control experiments, where we applied cortex buffer without oxytocin, correlations were unaffected (Fig. 3a, b). Next, we explored in more detail the correlations across the entire population of neurons and their changes after oxytocin application. We plotted correlations for all experiments during baseline against the correlations after oxytocin or cortex buffer application (Fig. 3c, d). We observed that oxytocin, but not cortex buffer applications, strongly decreased correlations across the entire population. There were essentially no neuronal pairs that showed increased correlations and the initial correlations did not appear to predict the oxytocin effect (Fig. 3c). Finally, we analyzed whether the distance between neurons determined the oxytocin effect and found that the oxytocin-mediated decrease in correlation increased with interneuronal distance (Fig. 3e, f), possibly reflecting increased lateral inhibition via SST^+^ interneurons (see below^27,28^).

**Figure 3.**
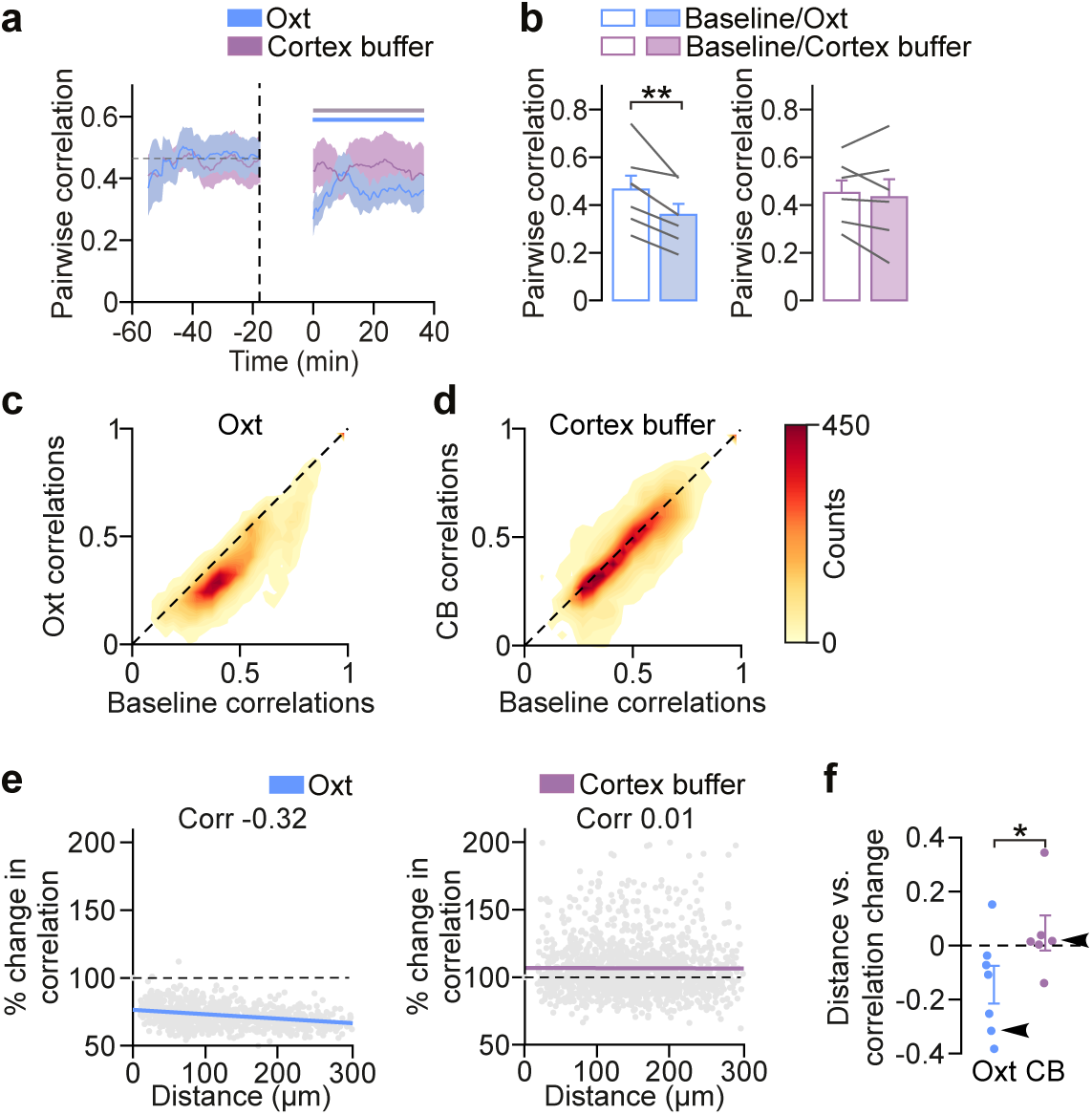
Oxytocin desynchronizes network activity in V1. **a.** Time-course of pairwise Pearson correlation coefficients before and after oxytocin and cortex buffer application. Dashed vertical line indicates the time point of oxytocin or cortex buffer application. Time course represents averages of seven-minute bins (for details see Materials and Methods). The horizontal bar indicates significant deviations from baseline as in Figure 1 (dark shades, paired two-tailed t-test, p < 0.05, without multi-measurement correction). **b.** Mean correlations after oxytocin or cortex buffer control applications. **p = 0.003 (n = 7 for oxytocin, n = 6 for cortex buffer, paired two-tailed t-test). **c.** Filled contour plot of the joint distribution of Pearson correlation coefficients. Pairwise correlations after oxytocin application plotted against baseline correlations. **d.** Pairwise correlations after cortex buffer (CB) plotted against baseline correlations. **e.** Change in mean pairwise correlations plotted against interneuronal distance for an example of oxytocin (left) and cortex buffer (right) conditions. Dashed lines indicate zero change in correlations. Colored lines indicate linear fits (left, p < 10^−10^; right, p = 0.47, paired two-tailed t-test). **f.** Pearson correlation coefficients between pairwise distances and percentages of change in correlations for all animals in oxytocin and cortex buffer condition. Arrowheads indicate the examples shown in **e**. *p = 0.037 (n = 7 for oxytocin, n = 6 for cortex buffer; unpaired two-tailed t-test).

Together, these findings demonstrated that oxytocin receptor activation modulates spontaneous activity patterns in a highly specific manner, since oxytocin affects spontaneous network activity in the visual, but not the somatosensory cortex. In addition, the frequency and the correlation of network events are specifically reduced after oxytocin application, leaving amplitude and duration unaffected.

### Oxytocin affects the E/I ratio differentially across sensory areas

To investigate whether the differential modulation of V1 and S1 by oxytocin can be explained by differences in the distribution of the oxytocin receptor in these areas, we used RNAscope to detect *Oxtr, Scl17a7* (coding for VGLUT1) and *Gad1* mRNA in the cortex at P10. We found that *Oxtr* is expressed in V1 as well as in S1 (Fig. 4). However, in V1 *Oxtr* co-localized almost exclusively with the *Gad1* signal (Fig. 4b, c), whereas in S1, the *Oxtr* signal co-localized with both the *VGLUT1* and *Gad1* signal (Fig. 4e, f). These observations suggested that differences between V1 and S1 in oxytocin receptor expression on the cellular level could explain oxytocin’s differential effect in these areas. To test this idea we examined how oxytocin regulated excitatory and inhibitory synaptic function in V1 and S1. Whole-cell patch-clamp recordings of layer 2/3 pyramidal neurons in slices from V1 showed that oxytocin affected neither the frequency (Fig. 5a, b) nor the amplitude of spontaneous excitatory postsynaptic currents (sEPSCs; Fig. 5c). Next, we measured spontaneous inhibitory postsynaptic currents (sIPSCs) at the reversal potential of glutamate receptor mediated currents. We found that the frequency of sIPSCs, in contrast to that of sEPSCs, was strongly increased after oxytocin bath application (Fig. 5d, e). The amplitude of sIPSCs was unaffected (Fig. 5f). To test whether this increase in frequency was mediated by the specific activation of the oxytocin and not the vasopressin 1A receptor^29^, which can be activated by oxytocin as well^30^, we blocked the oxytocin receptor using its specific antagonist OTA^31^. OTA prevented the oxytocin-mediated increase in sIPSC frequency entirely (Supplementary Fig. 1, fold-change oxytocin only: 4.95 ± 1.53; fold-change oxytocin + OTA: 0.98 ± 0.32, p = 0.0121, unpaired two-tailed Mann-Whitney test), demonstrating that this effect was mediated by the oxytocin receptor. Thus, activation of the oxytocin receptor increased the inhibitory tone in V1 dramatically, but did not affect excitatory synaptic transmission.

**Figure 4.**
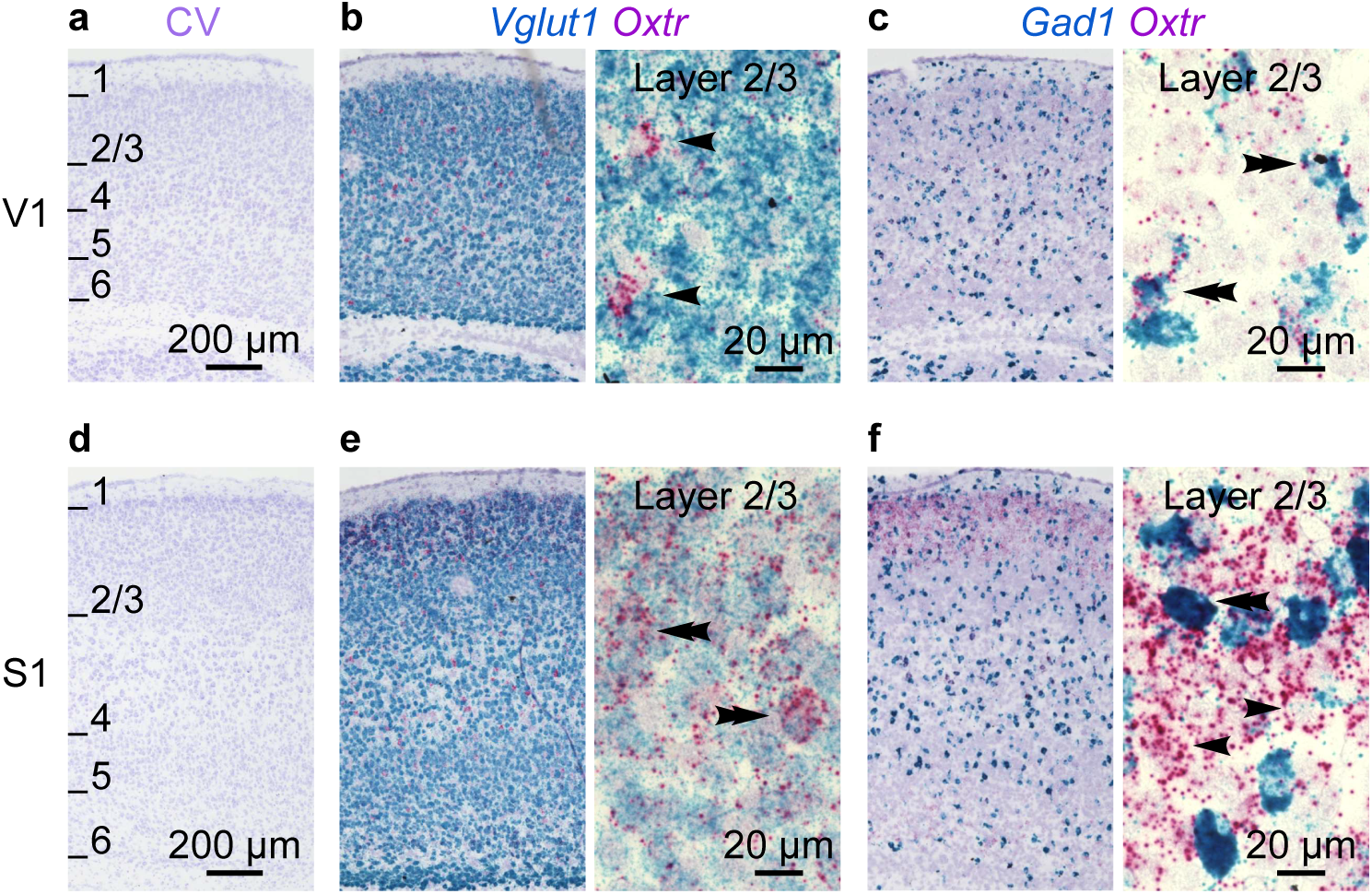
Oxytocin receptor mRNA is differentially expressed across sensory cortices. **a.** Cresyl violet Nissl staining of a sagittal section from V1 at low magnification. Numbers indicate cortical layers. **b.** Dual color RNAscope staining of *Vglut1* and O*xtr* mRNA in a V1 sagittal section. Left: low magnification. Right: high magnification. Note that the *Oxtr* signal was largely non-overlapping with the *Vglut1* signal (arrowheads indicate *Oxtr* positive/*Vglut1* negative neurons). **c.** Dual color RNAscope staining of *Gad1* and *Oxtr* mRNA in a V1 sagittal section. Left: low magnification. Right: high magnification, note that the *Oxtr* signal co-localized with the *Gad1* signal (double arrowhead). **d.** Cresyl violet Nissl staining of a sagittal section from S1 (same section as in **a**) at low magnification. Numbers indicate cortical layers. **e.** Dual color RNAscope staining of *Vglut1* and *Oxtr* mRNA in an S1 sagittal section. Left: low magnification. Right: high magnification, note the superposition of the two signals (double arrowheads). **f.** Dual color RNAscope staining of *Gad1* and *Oxtr* mRNA in an S1 sagittal section. Left: low magnification. Right: high magnification, note that the *Oxtr* signal co-localized only partially with the *Gad1* signal (arrowheads indicate *Oxtr* positive/*Gad1* negative neuron, double arrowhead indicates co-localization of both signals).

**Figure 5.**
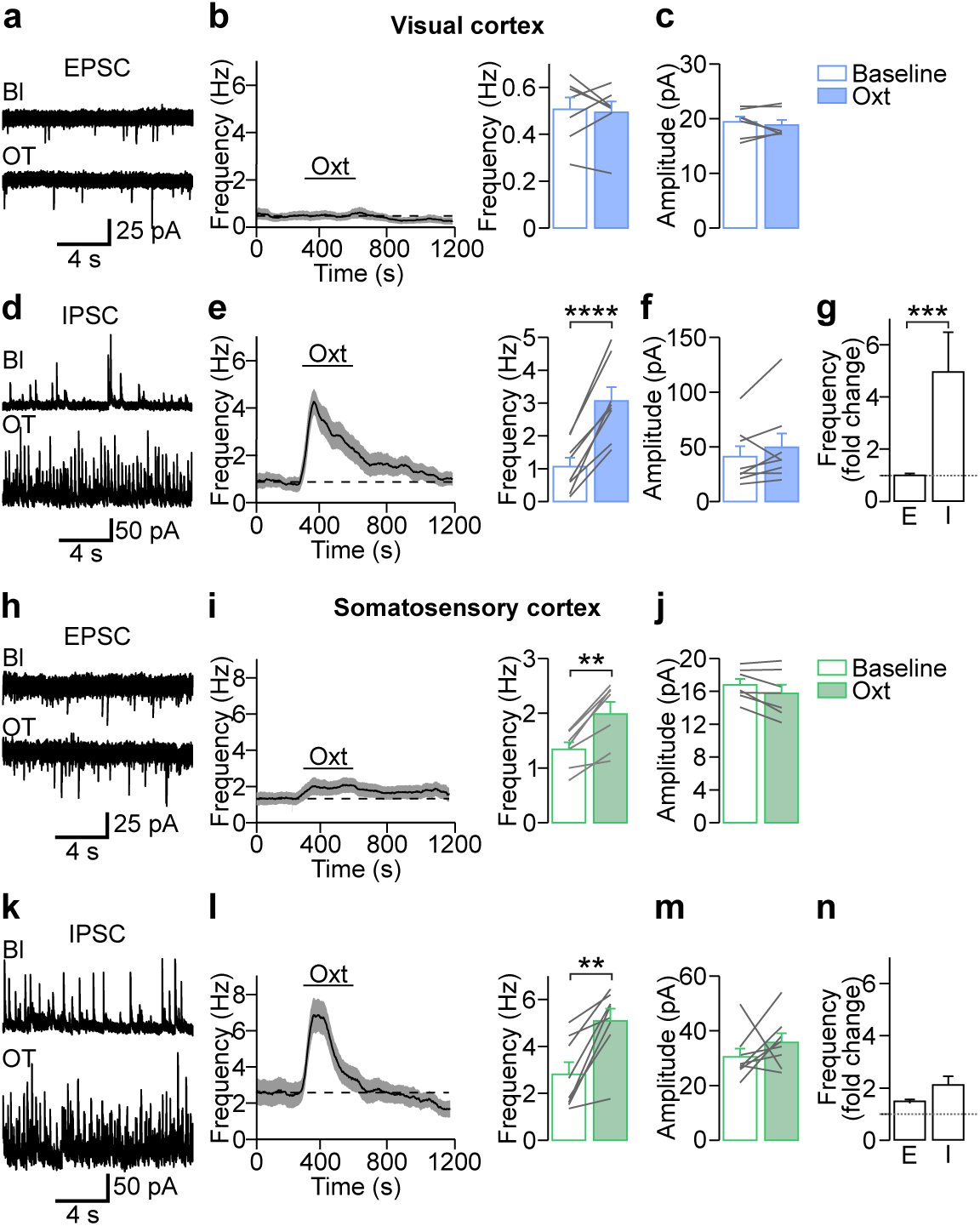
Oxytocin affects synaptic activity differentially across sensory cortices. **a.** Voltage-clamp recordings of spontaneous excitatory postsynaptic currents (sEPSCs) during baseline (Bl) and after oxytocin bath application in acute visual cortex slices. **b.** Frequency of sEPSCs before, during and after oxytocin application. In the visual cortex, oxytocin did not affect EPSC frequency. p > 0.5 (n = 7 cells, paired two-tailed t-test). **c.** Oxytocin did not affect sEPSC amplitude. p > 0.5 (n = 7 cells, paired two-tailed t-test). **d.** Spontaneous inhibitory postsynaptic currents (sIPSCs) before and after oxytocin application. **e.** Oxytocin led to a strong increase in sIPSCs. ****p = 8.21*10^−5^ (n = 8, paired two-tailed t-test). **f.** Oxytocin did not affect the amplitude of sIPSCs. p > 0.5 (n = 8 cells, paired two-tailed t-test). **g.** In the visual cortex, oxytocin led to a 5 times increase of sIPSCs, but sEPSCs were unaffected. ***p = 0.0003 (n = 7 and n = 8 cells for V1 sEPSCs and sIPSCs, respectively, Mann-Whitney test). **h.** Voltage-clamp recordings of sEPSCs before and after oxytocin bath application in somatosensory cortex slices. **i.** Application of oxytocin increased the frequency of sEPSCs in the somatosensory cortex slightly. **p = 0.0014 (n = 7, paired two-tailed t-test). **j.** Oxytocin did not affect the amplitude of sEPSCs. p > 0.5 (n = 7 cells, paired two-tailed t-test). **k.** sIPSCs before and after oxytocin application. **l.** Oxytocin led to a transient increase in sIPSCs. **p = 0.002 (n=8, paired two-tailed t-test). **m.** Oxytocin did not affect the amplitude of sIPSCs. p > 0.5 (n = 8 cells, paired two-tailed t-test). **n.** In the somatosensory cortex, oxytocin led to similar increases in sIPSC and sEPSC frequency. p = 0.1079 (n = 7 and n = 8 cells for S1 sEPSCs and sIPSCs, respectively, unpaired two-tailed t-test).

In contrast to V1, in S1 oxytocin bath application increased sEPSC frequency in layer 2/3 pyramidal cells (Fig. 5h, i). Again, sEPSC amplitude was unaffected (Fig. 5j). The frequency of sIPSCs was increased, but less pronounced than in V1 (Fig. 5k, l); amplitudes were unaffected (Fig. 5m). Thus, oxytocin shifted the E/I ratio in V1 towards inhibition (Fig. 5g, p < 0.001, two-ways ANOVA test), but did not affect E/I significantly in S1 (Fig. 5n). These experiments suggested that differences in the magnitude of inhibitory versus excitatory synaptic activity modulation accounted for the differences in the effect of oxytocin on spontaneous network activity in V1 versus S1.

### Oxytocin targets SST^+^ interneurons in the developing V1

Since we observed that oxytocin shaped spontaneous activity patterns strongly in V1, but not in S1, we focused next on the cellular mechanism of oxytocin mediated facilitation of inhibition in V1. First, we measured miniature IPSCs (mIPSCs) before and after oxytocin bath application and found that mIPSC frequency and amplitude were unaffected by oxytocin (Fig. 6a-c). This indicated that the number of inhibitory synapses, the density of postsynaptic receptors or changes in the presynaptic release machinery could not explain the increase in IPSC frequency described above.

**Figure 6.**
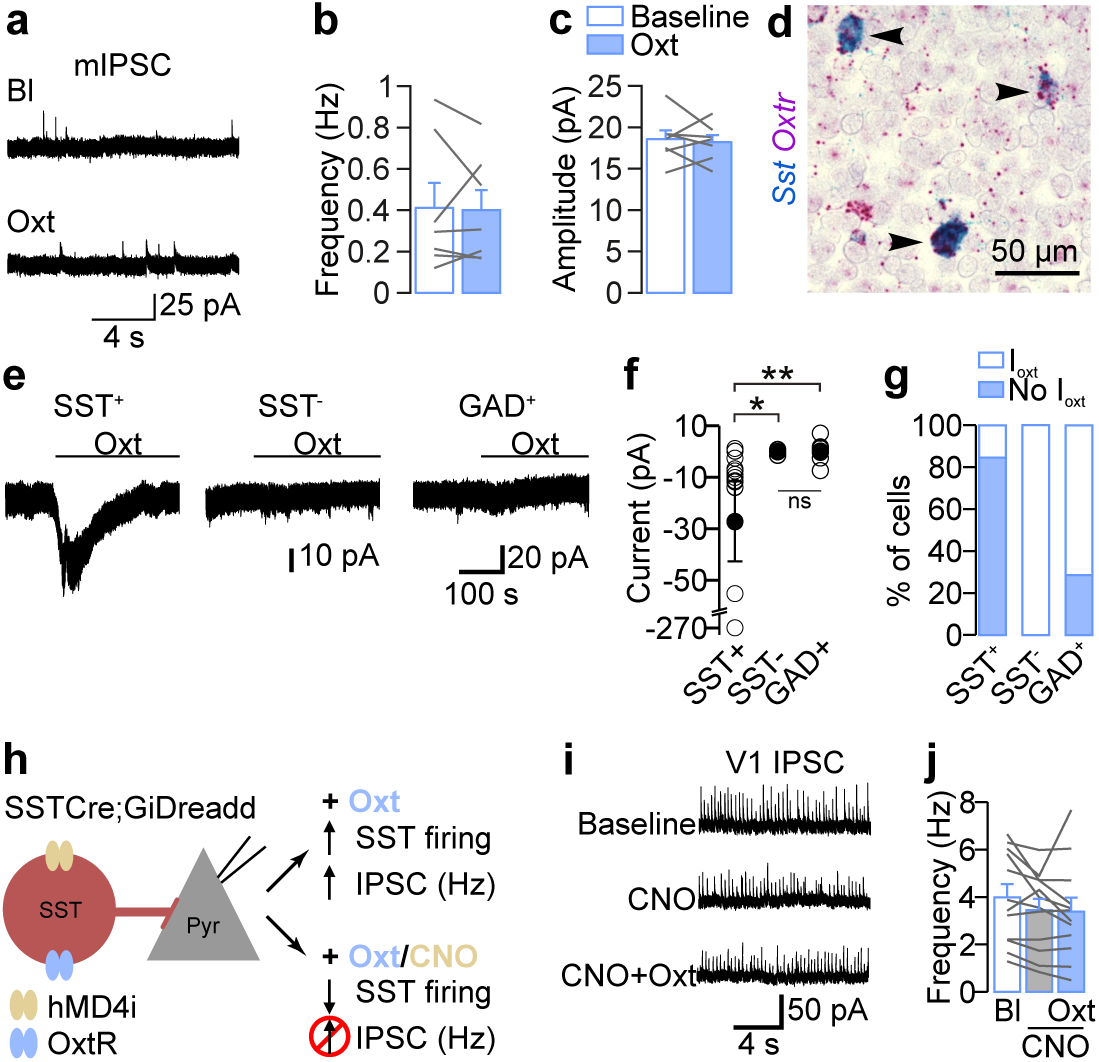
SST^+^ interneurons mediate the oxytocin-induced increase in inhibitory synaptic activity in V1. **a.** Voltage-clamp recordings of spontaneous miniature synaptic excitatory postsynaptic currents (mIPSCs) in the presence of tetrodotoxin (TTX, 10 µM) before and after oxytocin bath application in V1 slices. **b.** The frequency of mIPSCs was not affected by oxytocin. p > 0.5 (n = 7 cells, paired two-tailed t-test). **c.** The amplitude of mIPSCs was not affected by oxytocin. p > 0.5 (n = 7 cells, paired two-tailed t-test). **d.** V1 *Oxtr* and *Sst* mRNA transcripts. Note the superposition of the two signals (arrow head). **e.** Examples of voltage-clamp recordings at holding potential of −60 mV from a somatostatin (SST) expressing neuron (SST^+^, SST-Cre;Rosa26-TdTomato), an unlabeled neuron (SST^-^) and a GAD^+^ neuron (GAD^+^, GAD-Cre; Rosa26-TdTomato) before and after oxytocin application. Oxytocin induced an inward current in the SST^+^ neuron, but not in the unlabeled (most likely excitatory) neuron, nor in the GAD^+^ neuron. **f.** Group data of oxytocin-induced inward currents: SST^+^ n= 17 cells, SST^-^ n=7 cells and GAD^+^ n = 7 cells. *p = 0.0214, **p = 0.0044. Kruskal-Wallis test, followed by a Dunn test. **g.** Percentage of cells with oxytocin-induced inward currents. Almost all SST^+^ neurons show oxytocin-mediated currents, but none of the SST^-^ neurons and a fraction of GAD^+^ cells. **h.** Schematic representation of the experimental paradigm: pyramidal neurons in slices from a transgenic mouse expressing inhibitory DREADDs specifically in SST^+^ interneurons (SST-Cre;Rosa26-Gi-hMD4i) were patched in voltage-clamp mode. Oxytocin was applied while SST^+^ neuron activity was suppressed by CNO to test whether SST activation is required for the oxytocin induced increase in inhibitory synaptic activity. **i.** Example recordings of sIPSCs in baseline condition, in the presence of CNO alone and after bath application of oxytocin in the presence of CNO. **j.** Oxytocin did not increase sIPSC frequency in the presence of CNO. p > 0.05 (n = 12, one-way ANOVA).

To search for alternative causes of the increased inhibitory activity, we performed voltage-clamp recordings in inhibitory interneurons. We used two mouse lines expressing TdTomato to target inhibitory neurons in general (GAD2-Cre;Rosa26-TdTomato) or somatostatin expressing interneurons specifically (SST-Cre;Rosa26-TdTomato). We focused on SST^+^ interneurons for three reasons: first, RNA-seq data in adulthood showed that V1 oxytocin receptors are expressed primarily in SST^+^ interneurons (Supplementary Fig. 2a; Tasic et al., 2016, Allen Brain Atlas data portal: http://casestudies.brain-map.org/celltax)^32^. Second, we found here that already at P10 oxytocin receptor mRNA is expressed almost exclusively in SST^+^ interneurons in layer 2/3 of V1 (Fig. 6d). Finally, when we calculated the rate of rise (amplitude/rise time) of V1 IPSCs (Fig. 5d) we found that the rise rate was increased after oxytocin application (Supplementary Fig. 2b). Since the IPSC rise rate is larger for synapses that are located distally in the dendritic tree, this result indicated that oxytocin preferentially increased the activity of distal synapses. In fact, SST^+^ interneurons, in comparison with PV interneurons, target more distal dendrites of pyramidal cells^33^. Considering this evidence, we voltage-clamped SST^+^ neurons at −60 mV and blocked NMDA, AMPA and GABA_A_ receptor mediated currents using D-AP5, NBQX, and SR95531, respectively to prevent oxytocin-mediated network effects. In this configuration, we recorded oxytocin-mediated inward currents in almost all SST^+^ neurons (83%, amplitude: −27 ± 16 pA, Fig. 6e-g). Oxytocin did not trigger inward currents in any of the SST^-^ neurons in slices from the same SST-Cre;Rosa26-TdTomato mice. We concluded that oxytocin triggered depolarizing inward currents specifically in SST^+^ interneurons. We observed oxytocin induced inward currents in 29% of all GAD^+^ interneurons (Fig. 6g), in line with the proportion of SST^+^ neurons within the entire population of cortical interneurons^34^.

Together, these results indicated that oxytocin mediated the increase in inhibitory synaptic activity in V1 through activation of SST^+^ interneurons. To test this idea directly, we asked whether specifically downregulating SST^+^ interneurons might prevent the oxytocin-induced increase in overall inhibition (Fig. 6h). We performed voltage-clamp recordings of layer 2/3 neurons in acute slices from V1 of transgenic neonatal mice where SST^+^ interneurons expressed inhibitory designer receptors activated by designer drugs (iDREADDs, SSTCre;GiDreadd). Bath application of CNO, the activator of iDREADDs, did not affect the baseline frequency of sIPSCs (Fig. 6i, j), suggesting that SST^+^ interneurons were only sparsely active in our slice preparation. We then applied oxytocin and found that it did not increase sIPSC frequency in the presence of CNO (Fig. 6i, j) in contrast to our previous results in slices from wild type mice (Fig. 5e; fold-change WT: 4.95 ± 1.53, DREADD: 0.95 ± 0.07; p < 0.0001; Mann-Whitney test). Additional control experiments showed that oxytocin did increase the frequency of sIPSCs in neurons from iDREADD expressing animals in the absence of CNO (Supplementary Fig. 3a-c). These results showed that activation of SST^+^ interneurons is required for the oxytocin-induced increase in inhibition.

### Oxytocin enhances SST^+^ neuron excitability

Our results suggested that oxytocin triggered inward currents in SST^+^ interneurons (Fig. 6e-g) and that oxytocin-dependent enhancement of SST^+^ interneuron firing mediated its effect on network activity (Fig. 6h-j). Therefore, we investigated next whether and how oxytocin-induced inward currents enhanced SST^+^ neuron firing. In current-clamp mode, we injected a constant current to set the membrane potential of V1 layer 2/3 SST^+^ interneurons to −60 mV in the presence of the transmitter receptor blockers D-AP5, NBQX and SR95531. Then, we applied oxytocin while keeping the holding current constant. We observed that oxytocin induced a depolarization of 4.5 ± 0.4 mV (Fig. 7a), which exhibited the same transient temporal profile as the sIPSC frequency increase shown in Fig. 5e. Current-clamp recordings (Fig. 7b) revealed an increase in the firing rate of SST^+^ interneurons after oxytocin application (Fig. 7c). We further studied how oxytocin affected the action potential (AP) properties of SST^+^ interneurons. Oxytocin (1) depolarized the AP threshold slightly (+1.1 ± 0.4 mV; Fig. 7d), (2) increased the AP overshoot and amplitude (Fig. 7e, f, +2.01 ± 0.93 mV and +2.63 ± 0.78 mV, respectively), (3) broadened the AP width (+ 0.18 ± 0.03 ms at half-maximum; Fig. 7d, g) and (4) decreased the time required to generate an AP from the onset of current injection (−1.97 ± 0.32 ms; Fig. 7h). In addition, oxytocin modulated AP kinetics as it decreased the maximal speed of voltage change (dV/dt; Fig. 7i, j). Thus, oxytocin increased the firing capacity and AP properties of V1 layer 2/3 SST^+^ neurons, most likely by depolarizing their resting membrane potential, as described for PV interneurons in the hippocampus^35^.

**Figure 7.**
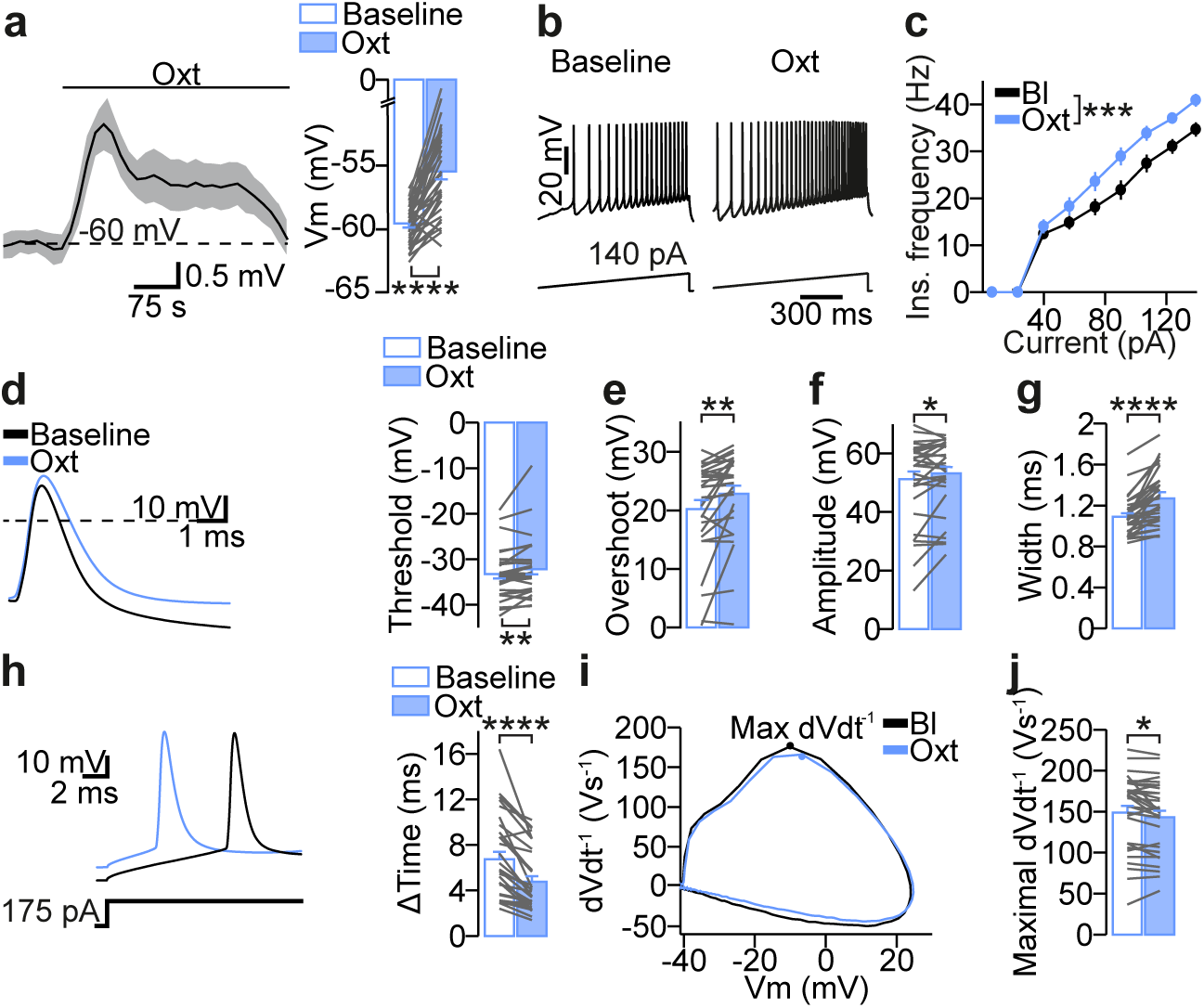
Oxytocin increases excitability of V1 SST^+^ interneurons. **a.** Current-clamp recording in an SST^+^ neuron before and after oxytocin bath application. Right, group data of SST^+^ interneuron membrane potential in baseline and oxytocin conditions. n = 34 cells (****p = 4.739*10^−10^, paired two-tailed t-test). **b.** Train of APs generated with a ramp protocol in current-clamp mode in baseline and oxytocin conditions. **c.** Instantaneous frequency versus current plot. ***p < 0.0001, two-ways ANOVA. **d.** Left, single action potentials aligned to the peak in baseline and oxytocin conditions. Right, group data of AP threshold in baseline and oxytocin conditions. **p = 0.0096, paired two-tailed t-test. **e.** AP overshoot. **p = 0.0032, paired two-tailed t-test. **f.** AP amplitude. *p =0.0375, paired two-tailed t-test. **g.** AP width. ****p = 1.833*10^−6^, paired two-tailed t-test. **h.** Left, single action potential in response to a square current step in baseline and oxytocin conditions. The time required to elicit an action potential was strongly reduced in the presence of oxytocin. ****p = 8.91*10^−7^, paired two-tailed t-test. **i.** Example action potential phase plot in baseline and oxytocin conditions. **j.** Maximal time derivative of membrane voltage in baseline and oxytocin conditions. *p = 0.0172, paired two-tailed t-test.

### Oxytocin modulates spontaneous activity patterns activating SST^+^ interneurons

Finally, we asked whether the oxytocin-mediated decrease in frequency of spontaneous network activity was the result of the activation of oxytocin receptors expressed in SST^+^ interneurons *in vivo*. We specifically inactivated SST^+^ interneurons by using a Cre-dependent inhibitory DREADD delivered by virus injections, which results in 80% of the SST^+^ interneurons expressing the hM4Di-DREADD construct^28^. Bath application of clozapine *in vitro* reduced the excitability of SST^+^ interneurons^28^, replicating previous findings that hM4Di-DREADD activation reduces the excitability of developing layer 2/3 neurons^36^. We performed *in vivo* wide-field calcium imaging to monitor spontaneous network activity in V1 and then activated the iDREADD receptor by injecting clozapine subcutaneously (Fig. 8a). Five minutes after clozapine injection, oxytocin was applied topically (Fig. 8a, b). In this condition oxytocin failed to decrease the frequency of spontaneous network events compared with oxytocin application alone (Fig. 8c, % of change clozapine + oxytocin: −18.1 ± 10.5; % of change oxytocin: −49.19 ± 6.2, p = 0.0231, unpaired two-tailed t-test). Area, amplitude and duration were not affected, either (Fig. 8d-f). Thus, oxytocin can modulate spontaneous activity patterns by specifically activating SST^+^ interneurons.

**Figure 8.**
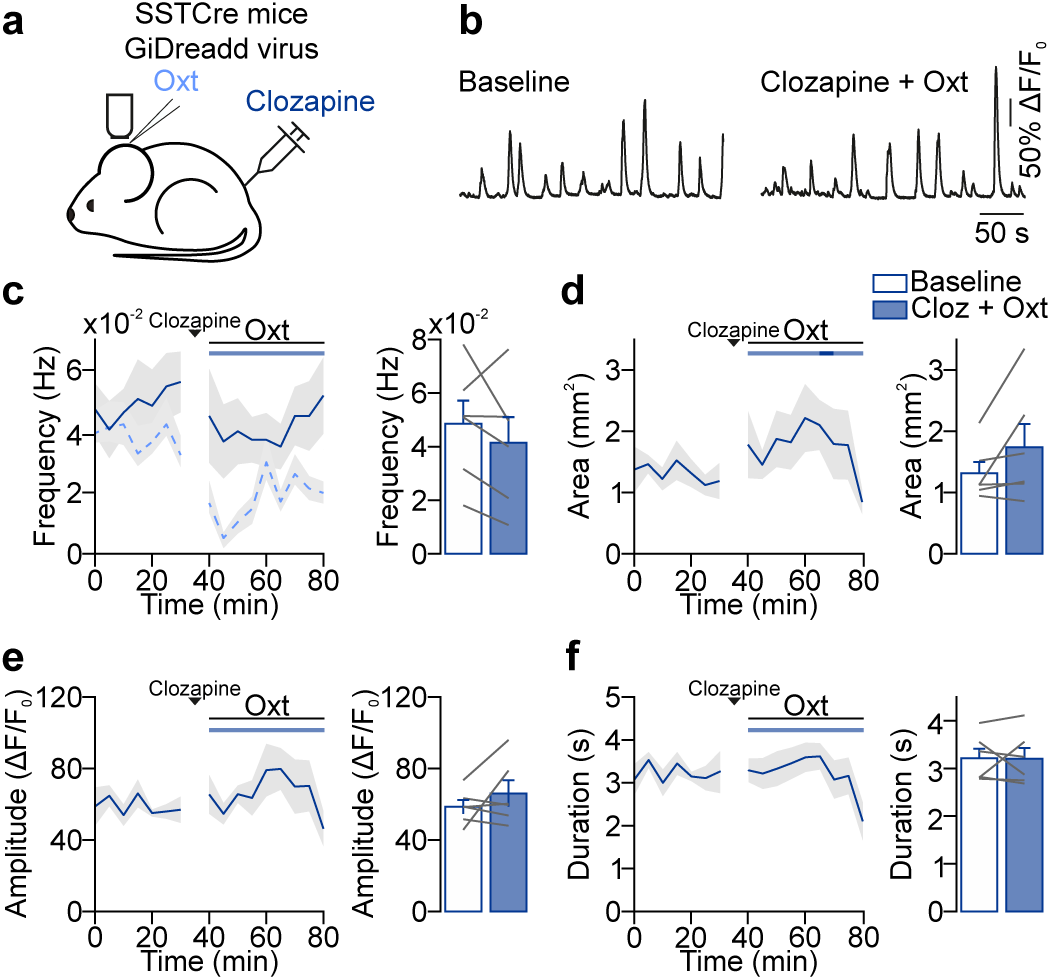
Oxytocin mediated decrease in network event frequency requires activation of SST^+^ interneurons *in vivo*. **a.** Schematic representation of the experimental design: wide-field calcium imaging of spontaneous network activity in SST-Cre mice where V1 neurons express GCaMP6 and GiDreadd after viral transduction. **b.** Fluorescent changes before and after clozapine + oxytocin application. **c.** Network event frequency during baseline and after oxytocin application. Time courses represent five minute-averages. The horizontal bars above the line plots indicate the time points when the values for each time bin did or did not differ significantly from baseline (dark and light shades, respectively, paired two-tailed t-test, p < 0.05, without multi-measurement correction). Dashed light blue curve represents data shown in Fig. 1**d**, V1, for comparison. **d.** Network event area. **e.** Network event amplitude. **f.** Network event duration.

## Discussion

In adults, oxytocin is a potent modulator of brain activity and behavior; however, its role during development has been less well established. Oxytocin is already present in the neonatal cortex and expression of the oxytocin receptor in the neocortex peaks during the first weeks of life^19–21^. Here, we demonstrate that before eye opening, during the second postnatal week, oxytocin modulates specific characteristics of spontaneous activity patterns in the visual cortex: it selectively increases SST^+^ interneuron excitability through activation of the oxytocin receptor and it sparsifies and decorrelates neuronal activity in V1 without affecting event amplitude, duration or spread across the cortex.

### The effect of oxytocin receptor activation differs between sensory modalities

A large body of evidence suggests that care behaviors of the mother induce activation of hypothalamic neurons and release of oxytocin in the pup’s brain; e.g. during mother-pup skin-to-skin contact^37^, anogenital stimulation^38^ or stroking stimuli^39^ and most likely after milk suckling activity^40^. How oxytocin reaches the developing cortex is not entirely clear. It might be released within V1, since hypothalamic oxytocin neurons project to the cortex including V1, at least in adult mice^18,21^. Alternatively, oxytocin may diffuse into the developing cortex after somatodendritic release from the hypothalamus into the third ventricle^19,29,40^. To fully disentangle how oxytocin reaches the developing cortex and to induce endogenous release, it will be required to adapt technical approaches currently available in adults^16,18,35^ to neonatal animals.

Independently of the source of oxytocin, we find here that this neuropeptide strongly decreases spontaneous network activity in V1, while its effect on spontaneous network activity in S1 is comparably mild. The differences between oxytocin’s effect on V1 and S1 network activity are consistent with differences in the cell-type expression of its receptor in these areas and their cellular responses to oxytocin. Oxytocin receptor mRNA expression co-localizes with the interneuron marker GAD1 in both V1 and S1, however its expression in excitatory neurons is higher in S1 than V1. Furthermore, oxytocin increases specifically inhibitory synaptic transmission in V1, but in S1 it results in a smaller and more balanced activation of both spontaneous inhibitory and excitatory currents. Thus, the specific effect of oxytocin on inhibitory signaling is most likely responsible for its effect on network activity patterns in V1. This conclusion is supported by our observation that the oxytocin receptor is transcribed and functional in SST^+^ interneurons and that pharmacogenetic suppression of SST^+^ neurons prevents the oxytocin-mediated decrease in network event frequency *in vivo*. Oxytocin increases inhibitory transmission also in various regions of the adult brain mainly acting on PV interneurons (e.g. in the piriform cortex, paraventricular nucleus (PVN), central amygdala and hippocampus^15,18,21,35,41^) or SST^+^ interneurons (e.g. in the auditory and prefrontal cortex ^16,17^).

The somatosensory system becomes responsive to sensory stimuli before the visual system. While vibrissal stimulation can elicit behavioral activity in pups at P3^42^ and their whiskers are necessary for nipple attachment to the mother and huddling by P5^42^, light-evoked responses are only present after P8^43^ and regular vision starts with eye opening. Therefore, our finding that oxytocin increases inhibitory tone and sparsifies network activity in V1, but not S1, during the second postnatal week may be related to the fact that V1 is less mature than S1 at this age. Physical contact with other pups or the mother in the nest triggers release of oxytocin in neonatal mice^37–39^. Thus, processing touch-mediated inputs in S1 may be important for immediate behavioral responses or early learning processes. In contrast, V1 is not yet required for behavioral control. Therefore, one might speculate that for example during social interactions in the nest, touch causes oxytocin release to sharpen tactile neuronal and behavioral responses by suppressing V1 spontaneous activity and preventing its spill over into S1^44^.

### SST^+^ interneuron activation accounts for the oxytocin-mediated increase in V1 inhibitory transmission

In the presence of synaptic transmission blockers, oxytocin induces an inward current (when held at −60 mV) in almost all SST^+^ interneurons and in a fraction of GAD^+^ interneurons, demonstrating that it activates a specific interneuron population in V1. Moreover, oxytocin-mediated depolarization of SST^+^ interneurons increases the frequency of inhibitory synaptic inputs in pyramidal cells five-fold and this increase is abolished entirely by SST^+^ interneuron inactivation. These observations are consistent with the presence of the oxytocin receptor mRNA in the young (Fig. 6d) and adult visual cortex in SST^+^ interneurons while it is hardly detected in other interneuron types, excitatory neurons or glia cells^32^. By contrast, in S1 both interneurons and excitatory neurons express the oxytocin receptor. Thus, the cellular distribution of the oxytocin receptor is brain region specific. Similar region-dependent differences in the cellular specificity of oxytocin receptor mediated responses have been observed in the hippocampus, where oxytocin receptors are activated in PV^+^ interneurons and pyramidal cells in CA2, but only in PV^+^ interneurons in CA1^15,35^.

Our finding that oxytocin increases inhibitory function through SST^+^ interneuron activation and decreases spontaneous activity during an important step in visual cortex development supports the idea that oxytocin modulation of inhibition facilitates the developmental progression across critical periods^45,46^. During development, changes in the E/I balance are important, because they determine the opening or closure of critical periods, during which sensory pathways become sensitive to experience^47^. Furthermore, the maturation of inhibition may be required for decreasing spontaneous activity^48^ and thereby increasing the relative importance of visually evoked activity. This relative increase in experience driven activity could initiate the critical period two weeks after eye opening^49^. Thus, oxytocin may be an important regulator of “phenotypic checkpoints” important for sensitive periods during postnatal development^50^.

### Oxytocin desynchronizes the network in a structured manner

Imaging activity across large populations of layer 2/3 neurons, we observed that oxytocin decreases pairwise correlations between neurons. This decorrelation is mediated most likely by oxytocin-induced activation of SST^+^ interneurons in agreement with previously published findings: activation of interneurons decreases neuronal correlations in general^51,52^. More specifically, lateral inhibition, a role attributed to SST^+^ interneurons, decorrelates spike trains^53^ or stimulus-evoked patterns^54^ and pharmacogenetic inactivation of SST^+^ interneurons increases pairwise correlations during spontaneous network activity in the developing visual cortex^28^. Our analyses suggest that correlations are down-regulated in a spatially specific manner: correlation decreases are more pronounced between pairs of relatively distant neurons. Neuronal correlations and the spatial extent of spontaneous network activity patterns determine their effectiveness in refining synaptic connections in the visual system^11,55,26^. Therefore, the regulation of spatial correlations through oxytocin may be necessary to shape spontaneous activity patterns to drive the refinement of synaptic connections and to prepare the emerging network optimally for computing visual inputs after eye opening. It might be tempting to speculate in this regard, that oxytocin contributes to improved visual acuity after tactile stimulation during development^56^.

During the second postnatal week, V1 activity patterns change from high correlation, high cell-participation patterns towards sparser and less correlated activity. Sparsification of activity patterns has been shown to be largely pre-programmed^57–59^. Since oxytocin signaling ramps up during the second postnatal week and sparsifies activity patterns as discussed above, oxytocin, and potentially other neuromodulators, may drive the sparsification and decorrelation seen in spontaneous activity patterns towards the onset of sensation. In this regard, it is interesting to note that activity patterns in the *fmr1* KO mouse, a model for the neurodevelopmental disorder fragile-X syndrome, show increased correlations in the developing somatosensory and visual cortices compared to wild-type littermates^60,61^. In addition, atypical oxytocin signaling has been implicated as a risk factor for neurodevelopmental disorders^62–64^. Thus, oxytocin may be important for healthy brain development to specifically shape activity patterns for synaptic refinement by regulating SST^+^ interneuron function.

## Experimental procedures

### Animals, *in utero* electroporation and surgery

All experimental procedures were approved by the institutional animal care and use committee of the Royal Netherlands Academy of Arts and Sciences. Neonatal C57BL/6J males and females from postnatal day 9 to 14 (P9-14) were used. *In utero* electroporation was performed as described previously^24^. Briefly, to perform calcium imaging of layer 2/3 pyramidal cells, GCaMP6s was cloned into pCAGGS (Addgene plasmid 40753^65^) and used in combination with DsRed in pCAGGS for visualization (gift from Christiaan Levelt). Pups were *in utero* electroporated at embryonic day (E) 16.5 after injection of the GCaMP6s (2 µg/µl) and DsRed (1 µg/µl) vectors into the ventricles. Electrode paddles were positioned to target the subventricular zone and 50 V pulses of 50 ms duration were applied.

For *in vivo* experiments, surgery for craniotomy was performed as described previously^7,28^. Briefly, pups were kept at 36-37 °C and anesthetized with 2% isoflurane and lidocaine was applied into the skin before neck muscle removal. A head bar was fixed above the V1/S1 region. Isoflurane was dropped to 0.7% before the imaging session. This lightly anesthetized state, which is characterized by rapid and shallow breathing and a relatively high heart rate, was maintained throughout the imaging session.

### Virus injection

Virus injections were performed at P0-1. pAAV-hSyn-DIO-hM4D(Gi)-mCherry and pAAV-hSyn-GCaMP6 plasmids were kindly packaged by Fred De Winter as described previously^28^. Neonatal mice were anesthetized by cold-induced hypothermia and kept cold in a stereotactic frame for pups (RWD Life Science). Stereotactic injections targeting V1 were performed with a microinjection pipet (Nanoject II, Drummond; volume 27 nl; mix of 1:1 AAV1-hSyn-DIO-hM4D(Gi)-mCherry and AAV1-hSyn-GCaMP6; from Bregma in mm: 0.3 posterior, 1.4 lateral). Immediately after injection, pups were kept warm on a heating pad and placed back to their mother after they awoke from anesthesia.

### Wide-field imaging

*In utero* electroporated or virus injected pups were used for calcium imaging of visual and somatosensory cortex. Calcium events were recorded with a Movable Objective Microscope (MOM, Sutter Instrument). Time-lapse recordings were acquired with a 4x objective (0.8 NA, Olympus) and blue light excitation from a Xenon Arc lamp (Lambda LS, Sutter Instrument Company). A CCD camera (Evolution QEi, QImaging) was controlled by custom-made LabVIEW (National Instruments) based software and images were acquired at a frame rate of 20 Hz.

Clozapine (Tocris) was injected subcutaneously (0.5 mg/kg) and oxytocin was applied five minutes after clozapine injection.

### 2-photon imaging

Bolus load of the calcium indicator Oregon Green 488 BAPTA-1 AM (OGB-1, Invitrogen) was performed as described^7^. Imaging was performed by using a two-photon microscope (MOM, Sutter, or A1RMP, Nikon) and a mode-locked Ti:Sapphire laser (MaiTai, Spectra Physics or Chamaleon, Coherent, λ = 810 nm). Consecutive xyt-stacks were acquired at a frame rate of 4-7 Hz through a 40x (0.8 NA, Olympus) or a 16x (0.8 NA, Nikon) water-immersion objective, controlled by ScanImage^66^ or NIS-Elements AR4.51.00 software (Nikon).

For *in vivo* experiments, oxytocin (1 µM, Sigma) was diluted in cortex buffer solution^7^ and applied topically at the craniotomy.

### Image analysis

Images were analyzed off-line with custom-written algorithms in Matlab (The MathWorks) as described previously^24^. Briefly, all recordings of an experiment were drift and motion corrected and aligned with respect to the first recording. ΔF/F_0_ stacks were generated using a moving average as F_0_ (window size 75 s). ROIs were hand-drawn using ImageJ (National Institutes of Health). For 2-photon calcium imaging experiments, events were first detected automatically in Matlab (The MathWorks) and then manually reviewed.

For pairwise correlation analysis, ΔF/F_0_ traces of five-minute long recording sessions were concatenated. The mean Pearson correlation coefficients were calculated for a moving seven-minute window. The filled contour plot was made plotting pairwise correlations after oxytocin (average of 40 minutes period after oxytocin) against baseline correlations (average of 40 minutes period before oxytocin).

For wide-field calcium imaging experiments, pixels below threshold were reduced to zero. The threshold was picked as the bending point of the cumulative histogram of all pixel-values across time. Network events were defined as groups of at least 300 non-zero pixels that were connected in time and/or space. These events were re-thresholded with a new threshold (the original threshold + 1.5 times the standard deviation of all pixels in the event). This last step was added after comparing the results of the automatic detection and manual detection, to make sure the two were consistent. The remaining non-zero pixels were considered as active.

### Patch-clamp experiments

Acute 300 µm coronal slices of the visual or somatosensory cortex were dissected between P9-P14. Pups were sacrificed by decapitation and their brains were immersed in ice-cold cutting solution (in mM): 2.5 KCl, 1.25 NaH_2_PO_4_, 26 NaHCO_3_, 20 Glucose, 215 Sucrose, 1 CaCl_2_, 7 MgCl_2_ (Sigma), pH 7.3-7.4, bubbled with 95%/5% O_2_/CO_2_. Slices were obtained with a vibratome (Microm HM 650V, Thermo Scientific) and subsequently incubated at 34°C in artificial cerebrospinal fluid (ACSF, in mM): 125 NaCl, 3.5 KCl, 1.25 NaH_2_PO_4_, 26 NaHCO_3_, 20 Glucose, 2 CaCl_2_, 1 MgCl2 (Sigma), pH 7.3-7.4. After 45 minutes, slices were transferred to the electrophysiology setup, kept at room temperature and bubbled with 95%/5% O_2_/CO_2_. For patch-clamp recordings, slices were transferred to a recording chamber and perfused (3 ml/min) with ACSF solution bubbled with 95%/5% O_2_/CO_2_ at 34°C.

Layer 2/3 pyramidal cells and interneurons were identified using an IR-DIC video microscope (Olympus BX51WI). GAD^+^ and SST^+^ interneurons were identified by the TdTomato protein fluorescence from the transgenic mice GAD2-Cre;Rosa26-TdTomato and SST-Cre;Rosa26-TdTomato, respectively. Quick change between bright-field imaging and epifluorescence was achieved using a foot-switch device^67^. Whole-cell voltage or current-clamp recordings were made with a MultiClamp 700B amplifier (Molecular Devices), filtered with a low pass Bessel filter at 10 kHz and digitized at 20-50 kHz (Digidata 1440A, Molecular Devices). Series resistance was assessed during recordings and neurons showing a series resistance >30 MΩ or a change >30% were discarded. Digitized data were analyzed offline using Clampfit 10 (Molecular Devices), Igor (WaveMetrics) and AxoGraph (Axograph Scientific).

Spontaneous and miniature IPSCs were recorded at a holding potential of 10 mV with glass pipettes (3-6 MΩ) containing (in mM): 115 CsCH_3_SO_3_, 10 Hepes, 20 CsCl, 2.5 MgCl_2_, 4 ATP disodium hydrate, 0.4 GTP sodium hydrate, 10 phosphocreatine disodium hydrate, 0.6 EGTA (Sigma), pH 7.3. For sIPSC recordings during selective silencing of SST^+^ interneurons, clozapine N-oxide (CNO) (10 µM, Tocris) was administered 1 minute prior to oxytocin (1 µM, Sigma). mIPSCs were recorded in the presence of tetrodotoxin (0.5 µM, Tocris). The oxytocin receptor antagonist desGly-NH2,d(CH2)5[D-Tyr2,Thr4]OVT (50 µM, synthesized and kindly donated by Dr Maurice Manning, University of Toledo) was applied for 5-10 minutes before oxytocin wash-in. sEPSCs were recorded at a holding potential of −60 mV (with junction potential correction) with an intracellular solution containing (in mM): 122 potassium gluconate, 10 Hepes, 13 KCl, 10 phosphocreatine disodium hydrate, 4 ATP magnesium salt, 0.3 GTP sodium hydrate (Sigma), pH 7.3.

### Analysis of electrophysiological experiments

m/sIPSCs and sEPSCs were detected using an Igor-based tool SpAcAn (Igor Pro 7, WaveMetrics). Frequency timelines of postsynaptic currents were built by calculating the frequency of 50-or 90-second bins. The 20-80 % rise time was calculated for each IPSC event and the rise rate was determined as amplitude/rise time (pA/ms). Oxytocin-induced currents in GAD^+^ and SST^+^ interneurons were recorded at a holding potential of −60 mV (with junction potential correction) in the presence of 10 µM SR95531, 10 µM NBQX, 50 µM D-AP5 (Tocris). Current-clamp recordings of SST^+^ interneurons were performed using the same KGluconate-based intracellular solution and in the presence of the synaptic blockers mentioned above. After breaking the seal, variable current injection was applied to keep the cells at −60 mV. The injected current was kept constant from the time of oxytocin wash-in. Excitability of SST^+^ interneurons was assessed with a one-step ramp protocol, from −100 pA to 140 pA at a rate of 96 pA/second. Single action potentials (APs) were elicited by injecting moderate pulses of current (<1 nA, <15 ms). Changes in membrane potential upon oxytocin treatment were calculated as the voltage difference between the trace exhibiting the peak effect and the last trace of baseline condition. A minority of cells that did not exhibit a depolarization (6%) were discarded for subsequent analyses. A voltage timeline was built by calculating the baseline membrane potential of 45-second bins. For analysis of SST^+^ interneuron excitability, the duration of the ramp was divided into 175-milisecond bins and the mean inter-spike interval (ISI) was calculated for each period bin. Then, the instantaneous frequency (ISI^-1^) was plotted against the mean current injected within the same bin. AP features were determined as follows: (1) overshoot was quantified as the amplitude of the AP above 0 mV. (2) Threshold was defined as the onset voltage with a slope greater than 50 mV/ms. (3) Amplitude was calculated as the voltage difference between the peak of the AP and its threshold. (4) Width was determined at half the amplitude of the AP. (5) ΔTime was described as the time difference between the onset of pulse injection and the time point when the membrane potential reached the action potential threshold. (6) dVdt^-1^ was defined as the first derivative of the voltage trace with respect to time.

### RNAscope

Animal protocols were approved by the Institutional Animal Care and Use Committee at Florida State University in accordance with state and federal guidelines (Guide for the Care and Use of Laboratory Animals of the National Institutes of Health). Fresh frozen brain tissue from C57BL/6J mice was sectioned in the sagittal plane at 20 μm in 6 series on SuperFrost Plus microscope slides and stored at −80°C until further processing. RNA transcripts were detected with RNAscope 2.5HD Duplex Assay (Cat. No. 322430, Advanced Cell Diagnostics (ACD), Hayward, CA). Synthetic oligonucleotide probes complementary to the nucleotide sequence 1198 – 2221 of *Oxtr* (NM_001081147.1; ACD Cat. No. 411101-C2), 464 - 1415 of *Slc17a7* (VGLUT1; NM_182993.2; ACD Cat. No. 416631), 62 – 3113 of *Gad1* (NM_008077.4; ACD Cat. No. 400951) and 18 – 407 of *Sst* (NM_009215.1; ACD Cat. No. 404631) were used. Slides were fixed for 2 hours in ice cold 4% paraformaldehyde (pH 9.5) followed by increasing concentrations of ethanol and dehydration in 100% ethanol overnight at −20°C. Slides were air dried for 10 minutes and boiled for 5 minutes in a target retrieval solution (ref. 322001, ACD), followed by 2 room temperature water rinses and a rinse in 100% ethanol. Slides were air dried, after which targeted sections were incubated with Protease Plus solution (ref. 322331, ACD) for 15 minutes at 40°C, followed by room temperature water rinses. These prepared slides were then probed for 2 hours with individual probe mixtures (*Oxtr* in the red channel 2 and the other probes in the blue-green channel 1) at 40°C. Unbound probes were rinsed off in wash buffer and slides were stored overnight in 5X SSC at room temperature. Signal amplification and detection were performed using the detailed instructions provided in the RNAscope 2.5HD Duplex Assay. Sections were counterstained with Gill’s hematoxylin (American Mastertech Scientific, Inc. Lodi, CA) and cover slipped with Vectamount (Vector Laboratories, Inc. Burlingame, CA). Images were captured with brightfield microscopy (Keyence BZ-X710, Keyence Corp., Osaka, Japan).

### Statistics

All data are shown as mean ± SEM. The number of animals and the test used for each analysis is specified in the Results section. To determine statistical differences we used Prism 7 (GraphPad). Sets of data ≥ 6 were tested for normality with a Shapiro-Wilk test, then a paired or unpaired t-test was applied for two-group comparisons. Comparisons between more than two groups were performed with one or two-way ANOVAs. For not normally distributed data or data < 6, the non-parametric Wilcoxon and Mann-Whitney tests were applied for paired or unpaired experiments, respectively, for two-group comparisons. Data sets with more than two groups were analyzed using the Kruskal-Wallis test.

## Author Contributions

Conceptualization: P.P.M and C.L; Experiments: P.P.M, A.N and E.H; Analysis: P.P.M, A.N and J.K., J.G.; Writing: P.P.M and C.L.

## Acknowledgments

We thank Corette Wierenga, Nicole Ropert and Marcus Howlett for critically reading this manuscript. Gertjan Houwen for his help with Matlab coding. Maurice Manning for the donation of the oxytocin receptor antagonist. Fred de Winter for the production of AAV-hSyn-DIO-hM4D(Gi)-mCherry and AAV-hSyn-GCaMP6 viruses. Marko Popovic for sharing with us the Rapid LED Switching System for one-photon microscopy. We thank the Helmut Kessels lab for sharing equipment and the Allen Brain Institute for permission to reproduce their data in Supplemental Figure 2. This work was supported by grants of the Netherlands Organization for Scientific Research (NWO, ALW Open Program grants, no. 819.02.017, 822.02.006 and ALWOP.216; ALW Vici, no. 865.12.001), the “Stichting Vrienden van het Herseninstituut” (all C. L.) and the Max Planck Society and the European Research Council StG 804824 (JG).

## References

1. Katz, L. C. & Shatz, C. J. Synaptic activity and the construction of cortical circuits. Science 274, 1133–1138 (1996).

2. Sanes, J. R. & Yamagata, M. Many paths to synaptic specificity. Annu. Cell DevBiol 25, 161–195 (2009).

3. Kirkby, L. A., Sack, G. S., Firl, A. & Feller, M. B. A Role for Correlated Spontaneous Activity in the Assembly of Neural Circuits. Neuron 80, 1129–1144 (2013).

4. Meister, M., Wong, R. O. L., Baylor, D. A. & Shatz, C. J. Synchronous bursts of action potentials in ganglion cells of the developing mammalian retina. Science 252, 939–943 (1991).

5. Feller, M. B., Wellis, D. P., Stellwagen, D., Werblin, F. S. & Shatz, C. J. Requirement for cholinergic synaptic transmission in the propagation of spontaneous retinal waves. Science 272, 1182–1187 (1996).

6. Ackman, J. B., Burbridge, T. J. & Crair, M. C. Retinal waves coordinate patterned activity throughout the developing visual system. Nature 490, 219–225 (2012).

7. Siegel, F., Heimel, J. A., Peters, J. & Lohmann, C. Peripheral and central inputs shape network dynamics in the developing visual cortex in vivo. Curr. Biol. 22, 253–258 (2012).

8. Colonnese, M. T., Shen, J. & Murata, Y. Uncorrelated Neural Firing in Mouse Visual Cortex during Spontaneous Retinal Waves. Front. Cell. Neurosci. 11, 289 (2017).

9. Weliky, M. & Katz, L. C. Disruption of orientation tuning in visual cortex by artificially correlated neuronal activity. Nature 386, 680–685 (1997).

10. Cang, J. et al. Development of Precise Maps in Visual Cortex Requires Patterned Spontaneous Activity in the Retina. Neuron 48, 797–809 (2005).

11. Burbridge, T. J. et al. Visual Circuit Development Requires Patterned Activity Mediated by Retinal Acetylcholine Receptors. Neuron 84, 1049–1064 (2014).

12. Bargmann, C. I. Beyond the connectome: how neuromodulators shape neural circuits. BioEssays News Rev. Mol. Cell. Dev. Biol. 34, 458–465 (2012).

13. Kaczmarek Leonard K. L., Irwin, B. Neuromodulation: The Biochemical Control of Neuronal Excitability. (Oxford University Press, 1986).

14. Lee, S.-H. & Dan, Y. Neuromodulation of Brain States. Neuron 76, 209–222 (2012).

15. Owen, S. F. et al. Oxytocin enhances hippocampal spike transmission by modulating fastspiking interneurons. Nature 500, 458–462 (2013).

16. Marlin, B. J., Mitre, M., D’amour, J. A., Chao, M. V. & Froemke, R. C. Oxytocin enables maternal behaviour by balancing cortical inhibition. Nature 520, 499–504 (2015).

17. Nakajima, M., Görlich, A. & Heintz, N. Oxytocin modulates female sociosexual behavior through a specific class of prefrontal cortical interneurons. Cell 159, 295–305 (2014).

18. Knobloch, H. S. et al. Evoked Axonal Oxytocin Release in the Central Amygdala Attenuates Fear Response. Neuron 73, 553–566 (2012).

19. Zheng, J.-J. et al. Oxytocin mediates early experience–dependent cross-modal plasticity in the sensory cortices. Nat. Neurosci. 17, 391–399 (2014).

20. Hammock, E. & Levitt, P. Oxytocin receptor ligand binding in embryonic tissue and postnatal brain development of the C57BL/6J mouse. Front. Behav. Neurosci. 7, (2013).

21. Mitre, M. et al. A Distributed Network for Social Cognition Enriched for Oxytocin Receptors. J. Neurosci. 36, 2517–2535 (2016).

22. Tyzio, R. et al. Maternal Oxytocin Triggers a Transient Inhibitory Switch in GABA Signaling in the Fetal Brain During Delivery. Science 314, 1788–1792 (2006).

23. Tyzio, R. et al. Oxytocin-mediated GABA inhibition during delivery attenuates autism pathogenesis in rodent offspring. Science 343, 675–679 (2014).

24. Winnubst, J., Cheyne, J. E., Niculescu, D. & Lohmann, C. Spontaneous activity drives local synaptic plasticity in vivo. Neuron 87, 399–410 (2015).

25. Stosiek, C., Garaschuk, O., Holthoff, K. & Konnerth, A. In vivo two-photon calcium imaging of neuronal networks. Proc.Natl.Acad.Sci.U.S.A 100, 7319–7324 (2003).

26. Leighton, A. H. & Lohmann, C. The Wiring of Developing Sensory Circuits-From Patterned Spontaneous Activity to Synaptic Plasticity Mechanisms. Front. Neural Circuits 10, 71 (2016).

27. Adesnik, H., Bruns, W., Taniguchi, H., Huang, Z. J. & Scanziani, M. A neural circuit for spatial summation in visual cortex. Nature 490, 226–231 (2012).

28. Leighton, A. H. et al. Modular control of spontaneous activity patterns by inhibitory signaling in the developing visual cortex. Revis. (2019).

29. Chini, B., Verhage, M. & Grinevich, V. The Action Radius of Oxytocin Release in the Mammalian CNS: From Single Vesicles to Behavior. Trends Pharmacol. Sci. 38, 982–991 (2017).

30. Schorscher-Petcu, A. et al. Oxytocin-Induced Analgesia and Scratching Are Mediated by the Vasopressin-1A Receptor in the Mouse. J. Neurosci. 30, 8274–8284 (2010).

31. Manning, M. et al. Oxytocin and Vasopressin Agonists and Antagonists as Research Tools and Potential Therapeutics. J. Neuroendocrinol. 24, 609–628 (2012).

32. Tasic, B. et al. Adult mouse cortical cell taxonomy revealed by single cell transcriptomics. Nat. Neurosci. 19, 335–346 (2016).

33. van Versendaal, D. & Levelt, C. N. Inhibitory interneurons in visual cortical plasticity. Cell. Mol. Life Sci. CMLS 73, 3677–3691 (2016).

34. Lee, S., Hjerling-Leffler, J., Zagha, E., Fishell, G. & Rudy, B. The largest group of superficial neocortical GABAergic interneurons expresses ionotropic serotonin receptors. J. Neurosci. Off. J. Soc. Neurosci. 30, 16796–16808 (2010).

35. Tirko, N. N. et al. Oxytocin Transforms Firing Mode of CA2 Hippocampal Neurons. Neuron 100, 593–608.e3 (2018).

36. Naskar, S. et al. The development of synaptic transmission is time-locked to early social behaviors in rats. Nat. Commun. 10, 1195 (2019).

37. Kojima, S., Stewart, R. A., Demas, G. E. & Alberts, J. R. Maternal Contact Differentially Modulates Central and Peripheral Oxytocin in Rat Pups During a Brief Regime of Mother–Pup Interaction that Induces a Filial Huddling Preference. J. Neuroendocrinol. 24, 831–840 (2012).

38. Caba, M., Rovirosa, M. J. & Silver, R. Suckling and genital stroking induces Fos expression in hypothalamic oxytocinergic neurons of rabbit pups. Dev. Brain Res. 143, 119–128 (2003).

39. Okabe, S., Yoshida, M., Takayanagi, Y. & Onaka, T. Activation of hypothalamic oxytocin neurons following tactile stimuli in rats. Neurosci. Lett. 600, 22–27 (2015).

40. Grinevich, V., Desarménien, M. G., Chini, B., Tauber, M. & Muscatelli, F. Ontogenesis of oxytocin pathways in the mammalian brain: late maturation and psychosocial disorders. Front. Neuroanat. 8, (2015).

41. Mühlethaler, M., Charpak, S. & Dreifuss, J. J. Contrasting effects of neurohypophysial peptides on pyramidal and non-pyramidal neurones in the rat hippocampus. Brain Res. 308, 97–107 (1984).

42. Landers, M. S. & Sullivan, R. M. Vibrissae-Evoked Behavior and Conditioning before Functional Ontogeny of the Somatosensory Vibrissae Cortex. J. Neurosci. 19, 5131–5137 (1999).

43. Colonnese, M. T. et al. A Conserved Switch in Sensory Processing Prepares Developing Neocortex for Vision. Neuron 67, 480–498 (2010).

44. Ackman, J. B., Zeng, H. & Crair, M. C. Structured dynamics of neural activity across developing neocortex. bioRxiv 012237 (2014) doi:10.1101/012237.

45. Grinevich, V. & Stoop, R. Interplay between Oxytocin and Sensory Systems in the Orchestration of Socio-Emotional Behaviors. Neuron 99, 887–904 (2018).

46. Hammock, E. A. D. Developmental Perspectives on Oxytocin and Vasopressin. Neuropsychopharmacology 40, 24–42 (2015).

47. Hensch, T. K. CRITICAL PERIOD PLASTICITY IN LOCAL CORTICAL CIRCUITS. Nat. Rev. Neurosci. 6, 877–888 (2005).

48. Colonnese, M. T. Rapid Developmental Emergence of Stable Depolarization during Wakefulness by Inhibitory Balancing of Cortical Network Excitability. J. Neurosci. 34, 5477–5485 (2014).

49. Toyoizumi, T. et al. A Theory of the Transition to Critical Period Plasticity: Inhibition Selectively Suppresses Spontaneous Activity. Neuron 80, 51–63 (2013).

50. Ben-Ari, Y. & Spitzer, N. C. Phenotypic checkpoints regulate neuronal development. Trends Neurosci. 33, 485–492 (2010).

51. Cardin, J. A. Inhibitory Interneurons Regulate Temporal Precision and Correlations in Cortical Circuits. Trends Neurosci. 41, 689–700 (2018).

52. Giridhar, S., Doiron, B. & Urban, N. N. Timescale-dependent shaping of correlation by olfactory bulb lateral inhibition. Proc. Natl. Acad. Sci. 108, 5843–5848 (2011).

53. Arevian, A. C., Kapoor, V. & Urban, N. N. Activity-dependent gating of lateral inhibition in the mouse olfactory bulb. Nat. Neurosci. 11, 80–87 (2008).

54. Friedrich, R. W. & Laurent, G. Dynamic optimization of odor representations by slow temporal patterning of mitral cell activity. Science 291, 889–894 (2001).

55. Xu, H.-P. et al. Spatial pattern of spontaneous retinal waves instructs retinotopic map refinement more than activity frequency. Dev. Neurobiol. 75, 621–640 (2015).

56. Guzzetta, A. et al. Massage Accelerates Brain Development and the Maturation of Visual Function. J. Neurosci. 29, 6042–6051 (2009).

57. Che, A. et al. Layer I Interneurons Sharpen Sensory Maps during Neonatal Development. Neuron 99, 98–116 (2018).

58. Golshani, P. et al. Internally Mediated Developmental Desynchronization of Neocortical Network Activity. J. Neurosci. 29, 10890–10899 (2009).

59. Rochefort, N. L. et al. Sparsification of neuronal activity in the visual cortex at eye-opening. Proc.Natl.Acad.Sci.U.S.A 106, 15049–15054 (2009).

60. Cheyne, J. E., Zabouri, N., Baddeley, D. & Lohmann, C. Spontaneous activity patterns are altered in the developing visual cortex of the Fmr1 knockout mouse. Front. Neural Circuits 13, (2019).

61. Goncalves, J. T., Anstey, J. E., Golshani, P. & Portera-Cailliau, C. Circuit level defects in the developing neocortex of Fragile X mice. Nat Neurosci 16, 903–909 (2013).

62. Hammock, E. A. D. & Young, L. J. Oxytocin, vasopressin and pair bonding: implications for autism. Philos. Trans. R. Soc. Lond. B. Biol. Sci. 361, 2187–2198 (2006).

63. Insel, T. R., O’Brien, D. J. & Leckman, J. F. Oxytocin, vasopressin, and autism: is there a connection? Biol. Psychiatry 45, 145–157 (1999).

64. Muscatelli, F., Desarménien, M. G., Matarazzo, V. & Grinevich, V. Oxytocin Signaling in the Early Life of Mammals: Link to Neurodevelopmental Disorders Associated with ASD. Curr. Top. Behav. Neurosci. 35, 239–268 (2018).

65. Chen, T.-W. et al. Ultrasensitive fluorescent proteins for imaging neuronal activity. Nature 499, 295–300 (2013).

66. Pologruto, T. A., Sabatini, B. L. & Svoboda, K. ScanImage: flexible software for operating laser scanning microscopes. BiomedEng Online 2, 13 (2003).

67. Battefeld, A., Popovic, M. A., van der Werf, D. & Kole, M. H. P. A Versatile and Open-Source Rapid LED Switching System for One-Photon Imaging and Photo-Activation. Front. Cell. Neurosci. 12, (2019).

