## Supplementary Information for "Oxytocin shapes spontaneous activity patterns in the developing visual cortex by activating somatostatin interneurons"

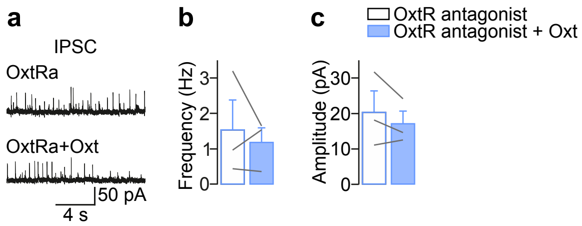


**Supplementary Figure 1.** The oxytocin receptor mediates the increase in sIPSC frequency after oxytocin application.

**a**. V1 sIPSCs in the presence of the oxytocin receptor antagonist (desGly-NH2,d(CH2)5[D-Tyr2,Thr4]OVT, donation from Maurice Manning, 50 µM) before (top) and after applying oxytocin (bottom).

**b**. When oxytocin receptors were blocked, oxytocin failed to increase the frequency of sIPSCs. N = 3 cells ($p > 0.05$, Wilcoxon test).

**c**. Amplitude of sIPSCs in the presence of the oxytocin receptor antagonist before and after oxytocin application. N = 3 cells ($p > 0.05$, Wilcoxon test).

**a**

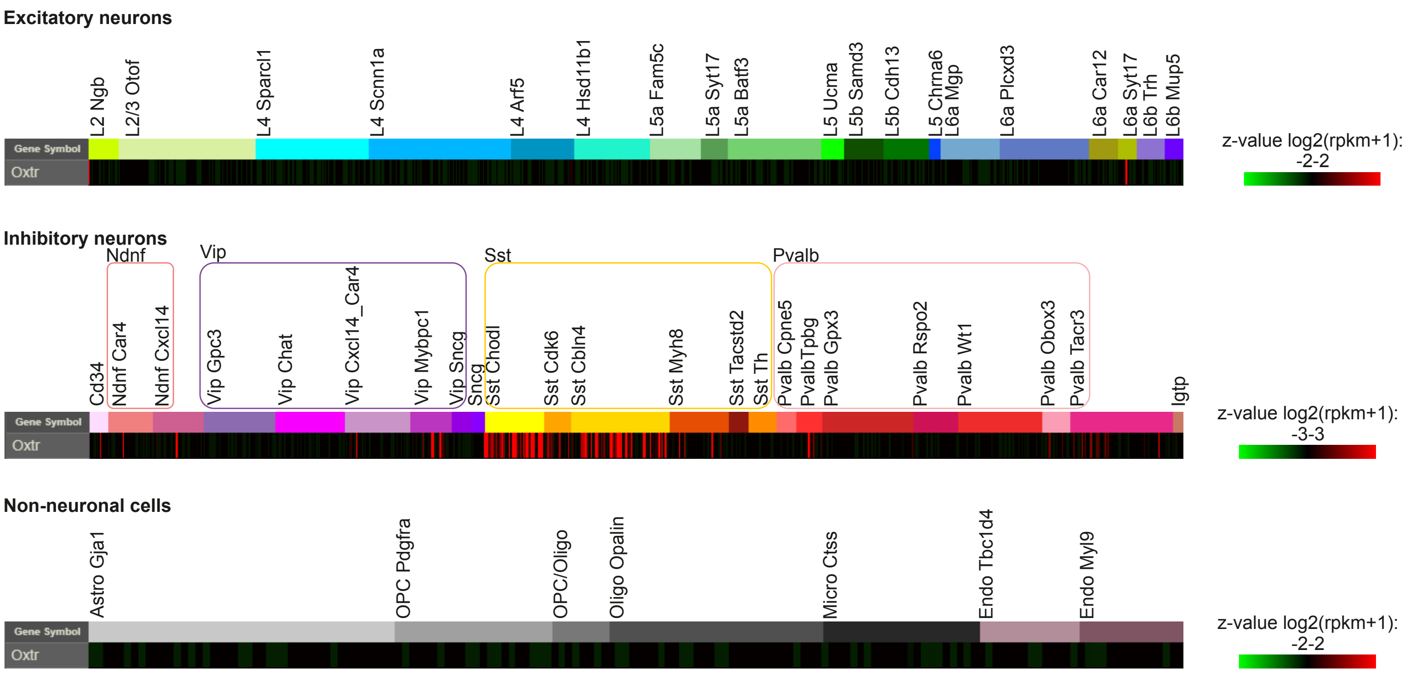


**Supplementary Figure 2a.** Adult visual cortex transcriptome.

**a.** Single cell RNA-sequencing of adult visual cortex for the oxytocin receptor gene (*Oxtr*). *Oxtr* is only expressed in interneurons, in particular in those of the somatostatin-expressing type. Adapted with permission from the Allen Brain Institute, Tasic et al., 2016, Allen Brain Atlas data portal: http://casestudies.brain-map.org/celltax.

**b**

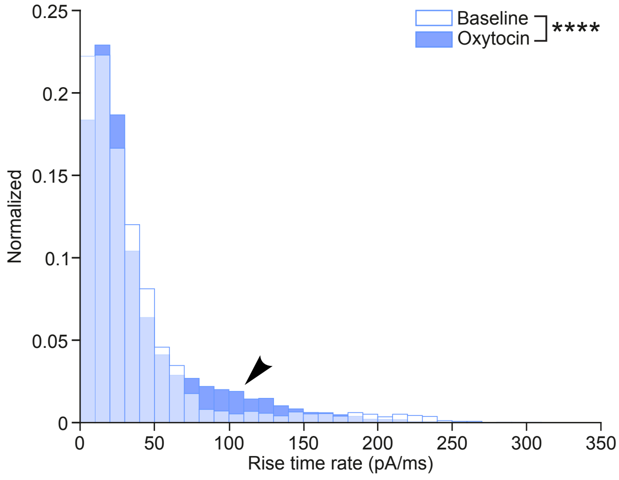


**Supplementary Figure 2b.** V1 sIPSC kinetics.

**b.** Rise time rate histogram of V1 sIPSCs. Oxytocin shifted the histogram to the left. N = 8 cells. Kolmogorov-Smirnov test, ****p < 0.0001.

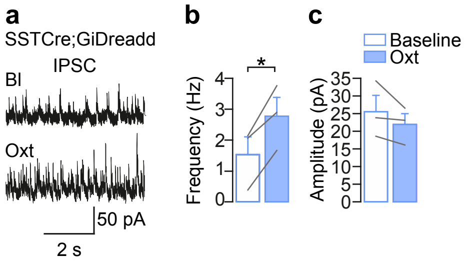


**Supplementary Figure 3.** Oxytocin increases the frequency of sIPSCs in SSTCre;GiDreadd mice.

**a**. sIPSCs before and after oxytocin application from a V1 layer2/3 pyramidal cell SSTCre;GiDreadd mouse in the absence of CNO.

**b**. sIPSC frequency. Oxytocin led to an increase in the frequency. N = 3 cells. paired two-tailed t-test, $*p < 0.05$.

**c**. Amplitudes of sIPSCs.
